# A role for fibroblast-derived SASP factors in the activation of pyroptotic cell death in mammary epithelial cells

**DOI:** 10.1101/2023.02.21.529458

**Authors:** Lisa M. Hom, Seunghoon Sun, Jamie Campbell, Pinyan Liu, Shannon Culbert, Ireland M. Murphy, Zachary T. Schafer

## Abstract

In normal tissue homeostasis, bidirectional communication between different cell types can shape numerous biological outcomes. Many studies have documented instances of reciprocal communication between fibroblasts and cancer cells that functionally change cancer cell behavior. However, less is known about how these heterotypic interactions shape epithelial cell function in the absence of oncogenic transformation. Furthermore, fibroblasts are prone to undergo senescence, which is typified by an irreversible cell cycle arrest. Senescent fibroblasts are also known to secrete various cytokines into the extracellular space; a phenomenon that is termed the senescence-associated secretory phenotype (SASP). While the role of fibroblast derived SASP factors on cancer cells has been well studied, the impact of these factors on normal epithelial cells remains poorly understood. We discovered that treatment of normal mammary epithelial cells with conditioned media (CM) from senescent fibroblasts (SASP CM) results in a caspase-dependent cell death. This capacity of SASP CM to cause cell death is maintained across multiple senescence-inducing stimuli. However, the activation of oncogenic signaling in mammary epithelial cells mitigates the ability of SASP CM to induce cell death. Despite the reliance of this cell death on caspase activation, we discovered that SASP CM does not cause cell death by the extrinsic or intrinsic apoptotic pathway. Instead, these cells die by an NLRP3, caspase-1, and gasdermin D (GSDMD)-dependent induction of pyroptosis. Taken together, our findings reveal that senescent fibroblasts can cause pyroptosis in neighboring mammary epithelial cells, which has implications for therapeutic strategies that perturb the behavior of senescent cells.

## Introduction

Heterotypic interactions between distinct cell types underlie a variety of diverse biological functions including shaping tissue architecture during normal development and maintaining tissue homeostasis (1–3). In addition, aberrant heterotypic signaling has been linked to a number of pathological conditions including neurodegenerative, cardiovascular, and autoimmune diseases (2,4–6). There has also been substantial research into heterotypic signaling in a variety of cancerous conditions (7–9). Heterotypic signaling during tumorigenesis often takes place between cells found in the tumor microenvironment (TME) and the cancer cells themselves (10–13). The TME is characterized by an array of cell types (e.g. fibroblasts, endothelial cells, immune cells) that are now appreciated to contribute significantly to tumor progression (14,15). While numerous studies have documented instances of reciprocal communication between fibroblasts and cancer cells that functionally change cancer cell behavior, much less is known about how these types of heterotypic interactions shape epithelial cell function in the absence of oncogenic transformation.

Relatedly, fibroblasts present in close proximity to epithelial cells have the propensity to undergo senescence, a process that is linked to aging and characterized by long-term loss of proliferative capacity (16–18). Senescent fibroblasts are well known to secrete inflammatory cytokines, which can have ramifications for cells in the local microenvironment. This phenomenon is characterized as the senescence associated secretory phenotype (SASP) and has been observed in response to multiple, distinct senescence inducers (19–21). Numerous lines of investigation have now demonstrated that the milieu of cytokines secreted by senescent fibroblasts can create a permissive environment for cancer cells (22–26). These data have motivated significant interest in the use of senolytic drugs, which specifically kill senescent cells, as a novel therapeutic strategy in multiple cancer types (27–29). However, the use of senolytic drugs to treat cancers is complicated by the fact that our understanding of how senescent fibroblasts can impact normal (non-cancerous) cells remains quite limited. For example, recent evidence suggests that senescent fibroblasts in the lung can sense injury-induced tissue inflammation, alter the stem cell niche, and facilitate the regeneration of epithelial barrier integrity (30). Therefore, additional studies that better clarify the relationship between senescent fibroblasts and non-cancerous epithelial cells are critically important to better understand the efficacy of this therapeutic strategy.

Here, we demonstrate that the induction of senescence (by distinct stimuli) in fibroblasts can result in the secretion of SASP factors that negatively impact the viability of normal mammary epithelial cells. In addition, the expression of activated oncogenes in normal mammary epithelial cells protects these cells from cell death caused by SASP factors. These findings suggest that cell death caused by SASP factors could functionally restrict extraneous growth of mammary epithelial cells prior to oncogenic transformation. When investigating the mechanism underlying cell death by SASP factors, we found that despite the necessity of caspase activation, this cell death was neither a consequence of extrinsic nor intrinsic-pathway mediated apoptosis. Instead, we found that SASP factors cause a pyroptotic cell death that is dependent on the NLRP3 inflammasome, caspase-1, and gasdermin D (GSDMD). Taken together, our findings reveal that senescent fibroblasts can cause pyroptosis in neighboring mammary epithelial cells, which has implications for therapeutic strategies that eliminate senescent cells.

## Results

### Conditioned media from senescent fibroblasts promotes caspase-dependent cell death in non-cancerous mammary epithelial cells

In order to study the effects of the senescence-associated secretory phenotype (SASP) in fibroblasts on the viability of non-cancerous mammary epithelial cells, we first induced senescence in BJ fibroblasts, which have been utilized extensively for studies examining fibroblast senescence (31–35). To do so, we treated BJ cells with the DNA-damaging agent bleomycin, a well-known inducer of senescence and SASP in this cell line (25,36). To confirm senescence induction in bleomycin treated cells, we measured senescence-associated-β-galactosidase activity (Figure 1A) and determined lamin B1 protein abundance (Figure 1B), which is known to decrease during senescence induction (37). Furthermore, we assessed the mRNA levels of IL-6 and IL-8 in bleomycin treated cells and confirmed that the expression of these factors, known to be associated with the SASP (38), are significantly elevated upon bleomycin treatment (Figure 1C).

**Figure 1:**
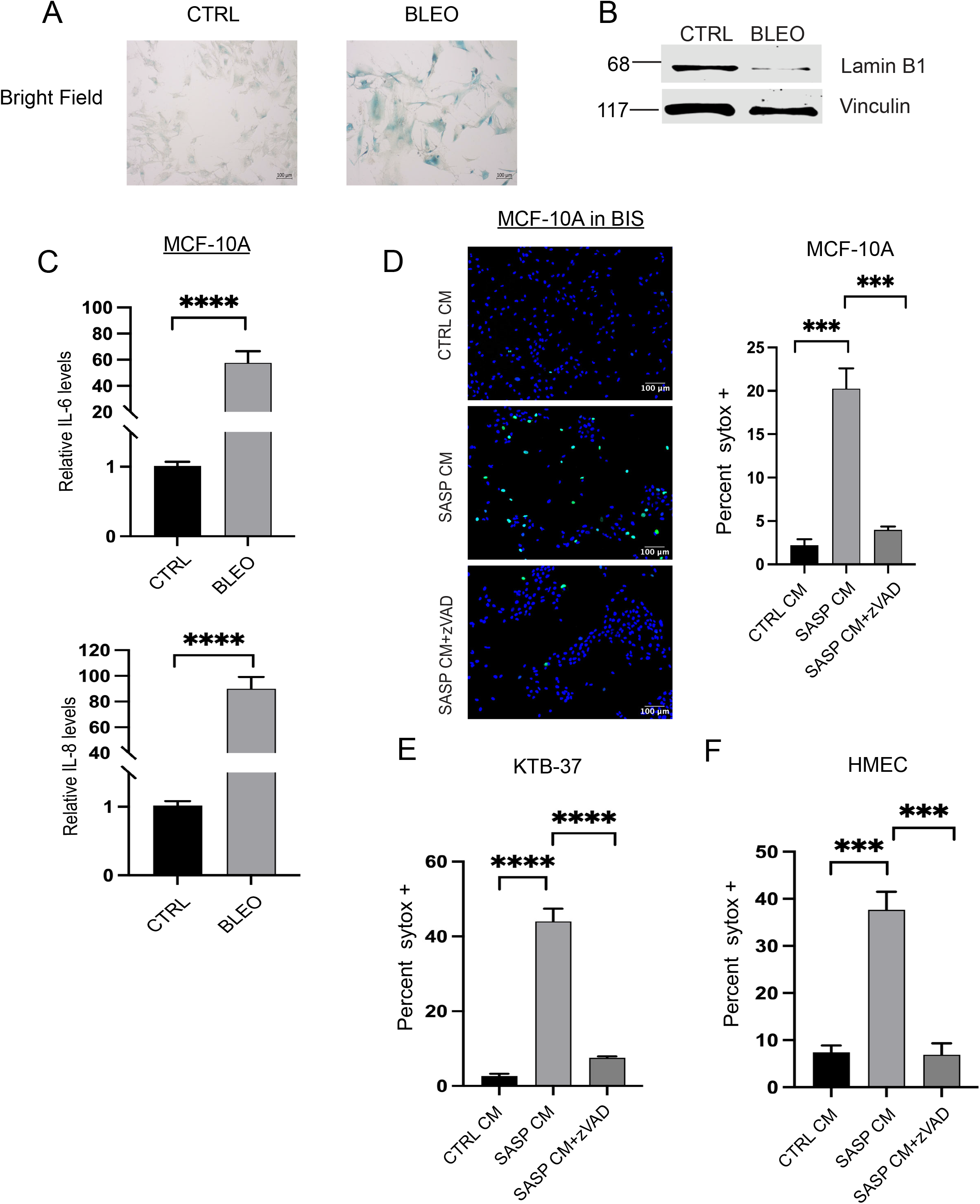
SASP CM from BIS fibroblasts causes cell death in normal mammary epithelial cells. **(A)** BJ fibroblasts were stained for Senescence-Associated β-Galactosidase (SA-β-Gal) activity to confirm the induction of senescence in BJ fibroblasts treated with DMSO or bleomycin. Images were taken at 10X in brightfield. Scale bar 100μm. **(B)** Cell lysates from BJ fibroblasts treated with DMSO or bleomycin were immunoblotted for lamin B1 protein levels as a marker for senescence induction. **(C)** qRT-PCR was performed to measure the relative mRNA levels of IL-6 and IL-8 in BJ fibroblasts treated with either DMSO or bleomycin to validate SASP induction. n=3 **(D)** (Left) Representative immunofluorescence images of MCF-10A cells treated with CTRL or SASP conditioned media (CM) from bleomycin induced senescent (BIS) fibroblasts and stained with Hoechst and SYTOX Green. Images were taken at 10X and scale bar is 100μm. z-VAD-fmk treatment was included to assess the impact of caspase activation on the abundance of SYTOX Green positive cells. (Right) Quantification of images represented as the percentage of SYTOX Green positive cells out of the total number of cells was depicted. **(E)** KTB-37 cells were treated with CTRL, SASP CM or SASP CM+z-VAD-fmk collected from BIS BJ fibroblasts. Quantification represented as the percentage of SYTOX Green positive cells out of the total number of cells was depicted. **(F)** HMEC cells were treated with CTRL, SASP CM or SASP CM+z-VAD-fmk collected from BIS BJ fibroblasts. Quantification represented as the percentage of SYTOX Green positive cells out of the total number of cells was depicted. Unpaired two-tail t-test was performed for qRT-PCR data where * p<0.05, ****p<.0001, ***p<.001. Data are presented as mean +/− SEM. SYTOX data was analyzed by one-way ANOVA followed by Tukey comparison test. Graphs are representative data collected from a minimum of three biological replicates.

Following confirmation of senescence induction by bleomycin, we collected conditioned media (CM) from control (non-senescent) or senescent fibroblasts (hereafter referred to as CTRL CM or SASP CM, respectively) and assessed cell death in MCF-10A cells treated with each type of CM by SYTOX Green staining. Interestingly, treatment with SASP CM resulted in a significant induction of cell death in MCF-10A cells (Figure 1D). Moreover, SASP CM-induced cell death in MCF-10A cells was blocked by treatment with z-VAD-fmk (a pan-caspase inhibitor), suggesting that the mechanism of cell death is dependent on caspase activation (Figure 1D). To assess whether the induction of cell death by SASP CM was specific to MCF-10A cells, we expanded our studies to additional mammary epithelial cell lines. Indeed, SASP CM treatment resulted in robust cell death in both KTB-37 (Figure 1E) and HMEC cells (Figure 1F), which was averted by treatment with z-VAD-fmk.

To complement these studies, we next evaluated whether SASP CM could promote cell death when senescence was induced in fibroblasts through an alternative mechanism. We engineered BJ fibroblasts to undergo oncogene-induced senescence (OIS) through overexpression of the oncogene HRas (G12V) and confirmed senescence induction through senescence-associated-β-galactosidase activity (Figure 2A), decrease in lamin B1 (Figure 2B), and expression of SASP genes (Figure 2C). Indeed, we found that SASP CM collected from OIS fibroblasts also stimulated the induction of cell death in MCF-10A (Figure 2D), KTB-37 (Figure 2E), and HMEC (Figure 2F) cells, and in each case, the SASP CM-mediated cell death was abrogated by treatment with z-VAD-fmk. Taken together, these data suggest that senescence induction (by distinct mechanisms) can cause fibroblasts to secrete factors that provoke cell death in normal mammary epithelial cells.

**Figure 2:**
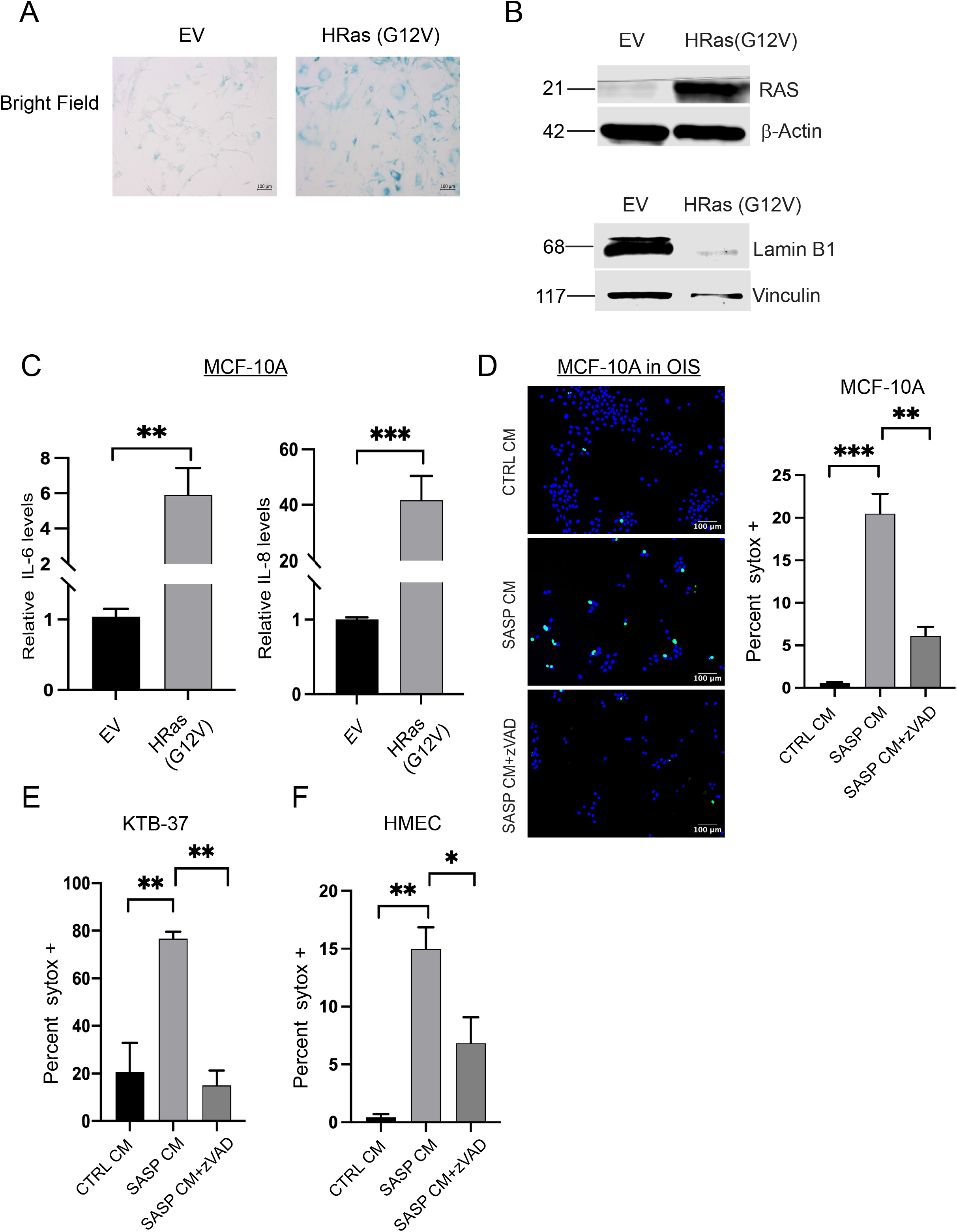
SASP CM from OIS fibroblasts causes cell death in normal mammary epithelial cells. **(A)** BJ fibroblasts were stained for SA-β-Gal activity to confirm the induction of senescence in BJ fibroblasts engineered to undergo oncogene-induced senescence (OIS) through overexpression of HRas (G12V). Empty vector (EV) control is also included. Representative images were taken at 10X in brightfield and scale bar is 100μm. **(B)** Cell lysates from BJ fibroblasts with EV or HRas (G12V) were immunoblotted for Ras or lamin B1 protein levels as a marker for senescence induction. **(C)** qRT-PCR was performed to measure the relative mRNA levels of IL-6 and IL-8 in BJ fibroblasts engineered to overexpress EV or HRas (G12V). n=3 **(D)** Representative immunofluorescence images of MCF-10A cells treated with CTRL or SASP conditioned media (CM) from OIS fibroblasts stained with Hoechst and SYTOX green. Images were taken at 10X and the scale bar is 100μm. Quantification of images represented as the percentage of SYTOX Green positive cells out of the total number of cells was depicted. **(E)** KTB-37 cells treated with CTRL or SASP conditioned media (CM) from OIS were stained with Hoechst and SYTOX Green. Quantification of images represented as the percentage of SYTOX Green positive cells out of the total number of cells was depicted. **(F)** HMEC cells treated with CTRL or SASP conditioned media (CM) from OIS stained with Hoechst and SYTOX Green. Quantification of images represented as the percentage of SYTOX Green positive cells out of the total number of cells was depicted. Unpaired two-tail t-test was performed for qRT-PCR data where ns is no statistical significance, * p<0.05, ** p<0.01, ****p<.0001 and ***p<.001. Data are presented as mean +/− SEM. Sytox data was analyzed by one-way ANOVA followed by Tukey comparison test. Graphs are representative data collected from a minimum of three biological replicates.

### Oncogenic signaling desensitizes breast cancer cells to SASP CM-induced cell death

Given our results demonstrating that SASP CM can cause cell death in non-cancerous mammary epithelial cells, we questioned whether breast cancer cells would respond similarly to exposure to SASP CM. This is a particularly important factor to consider given that senescent fibroblasts are well-known constituents of the breast tumor microenvironment (39–41). To assess the capacity of senescent fibroblasts to secrete factors that alter cell death in breast cancer cells, we treated MDA-MB-231 (Figure 3A) or MDA-MB-436 (Figure 3B) cells with SASP CM and measured cell death. Using SASP CM from both bleomycin-induced senescence (BIS) and OIS, we found that SASP CM treatment did not induce cell death in either cell line (Figures 3A, 3B). We next sought to extend these studies in an isogenic background using MCF-10A cells, which we previously demonstrated were sensitive to cell death caused by SASP CM (Figures 1D, 2D). To do so, we stably expressed HRas (G12V), ErbB2, or an empty vector (EV) control in MCF-10A cells and confirmed successful expression by immunoblotting (Figures 3C, 3E). We then exposed MCF-10A cells expressing these oncogenic mutations (or the EV control) to SASP CM (collected after induction of BIS or OIS) and measured cell death. Indeed, expression of either HRas (G12V) (Figure 3D) or ErbB2 (Figure 3F) was sufficient to block the ability of SASP CM to induce cell death. Thus, while non-cancerous epithelial cells are sensitive to cell death caused by SASP CM, the activation of oncogenes in normal mammary epithelial cells prevent the induction of SASP CM-mediated cell death.

**Figure 3:**
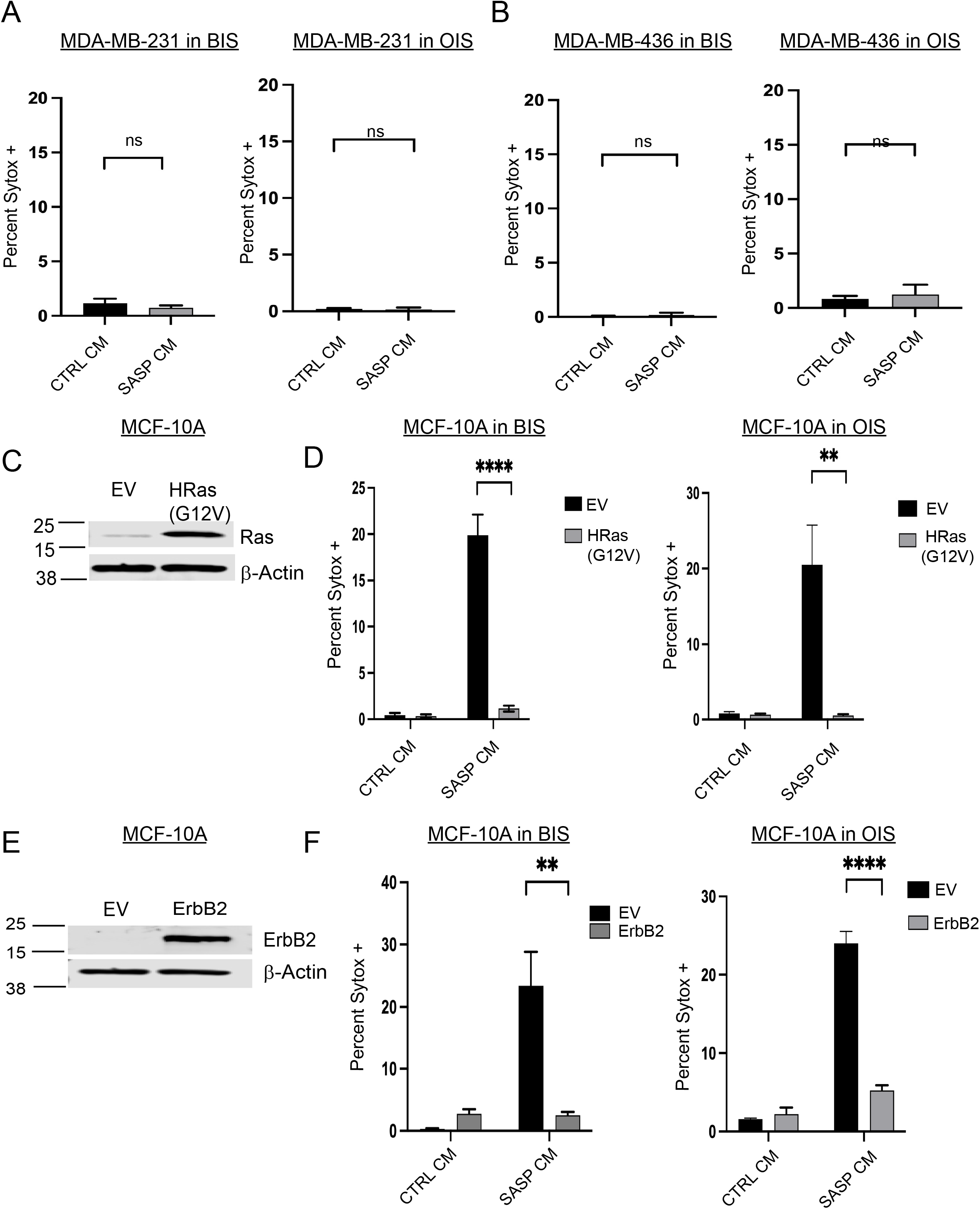
Breast cancer cells are resistant to SASP CM induced cell death. **(A,B)** MDA-MB-231 **(A)** and MDA-MB-436 **(B)** breast cancer cell lines were treated with CTRL or SASP CM collected from BIS or OIS fibroblasts and imaged using immunofluorescence microscopy. Cells were stained with Hoechst and SYTOX Green. Quantification of the percentage of SYTOX Green positive cells out of the total number of cells was depicted. **(C)** MCF-10A cells were engineered to overexpress either empty vector (EV) or the oncogene HRas (G12V). Overexpression was validated by immunoblot. **(D)** MCF-10A EV and HRas (G12V) overexpressing cells were treated with CTRL or SASP CM collected from BIS or OIS fibroblasts and imaged using immunofluorescence microscopy. Cells were stained with Hoechst and SYTOX Green. Quantification of the percentage of SYTOX Green positive cells out of the total number of cells was depicted. **(E)** MCF-10A cells were engineered to overexpress either EV or the oncogene ErbB2. Overexpression was validated by immunoblot. **(F)** MCF-10A EV and ErbB2 overexpressing cells were treated with CTRL or SASP CM collected from BIS or OIS fibroblasts and imaged using immunofluorescence microscopy. Cells were stained with Hoechst and SYTOX Green. Quantification of the percentage of SYTOX Green positive cells out of the total number of cells was depicted. SYTOX data were analyzed by oneway ANOVA followed by Tukey comparison test where ns is so statistical significance, * p<0.05, ** p<0.01, *** p<0.001 and **** p<0.0001. Data are presented as mean +/− SEM. Graphs are representative data collected from a minimum of three biological replicates. Isogenic manipulations were analyzed by two-way ANOVA followed by Tukey comparison test. Data are presented as mean +/− SEM. Graphs are representative data collected from a minimum of three biological replicates.

### SASP CM does not induce apoptotic cell death

We next sought to determine the molecular mechanism by which SASP-CM promotes cell death in mammary epithelial cells. Due to the fact that SASP CM-mediated cell death was blocked by the pan-caspase inhibitor z-VAD-fmk (Figures 1D–1F, 2D–2F), we posited that these cells were dying by apoptosis. We first investigated the role of the intrinsic apoptotic pathway, which is mediated by the release of cytochrome *c* from the mitochondria (42,43). To do so, we engineered MCF-10A cells to stably express Bcl-2, which blocks the release of cytochrome *c* from the mitochondria and thereby prevents downstream caspase activation (44). We confirmed Bcl-2 expression and function by immunoblotting for Bcl-2 and cleaved poly (ADP-ribose) polymerase (PARP) (Figure 4A). Interestingly, while Bcl-2 expression was able to effectively inhibit staurosporine-induced apoptosis, it was unable to block cell death caused by exposure of cells to SASP CM (from either BIS or OIS) (Figure 4B). Given these results, we next assessed the role of the extrinsic apoptotic pathway, which is characterized by ligand-mediated activation of death receptors (45,46). To do so, we utilized lentiviral-mediated transduction of shRNA in MCF-10A cells to lower the abundance of caspase-8 (Figure 4C), a key mediator of the extrinsic apoptotic pathway. Despite a substantial (shRNA-mediated) decrease in the abundance of caspase-8 protein, MCF-10A cells were still sensitive to cell death induced by SASP CM (from either BIS or OIS) (Figure 4D). Collectively, these data suggest that neither the intrinsic or extrinsic pathways of apoptosis mediate cell death caused by SASP CM, indicating the cell death is likely non-apoptotic.

**Figure 4:**
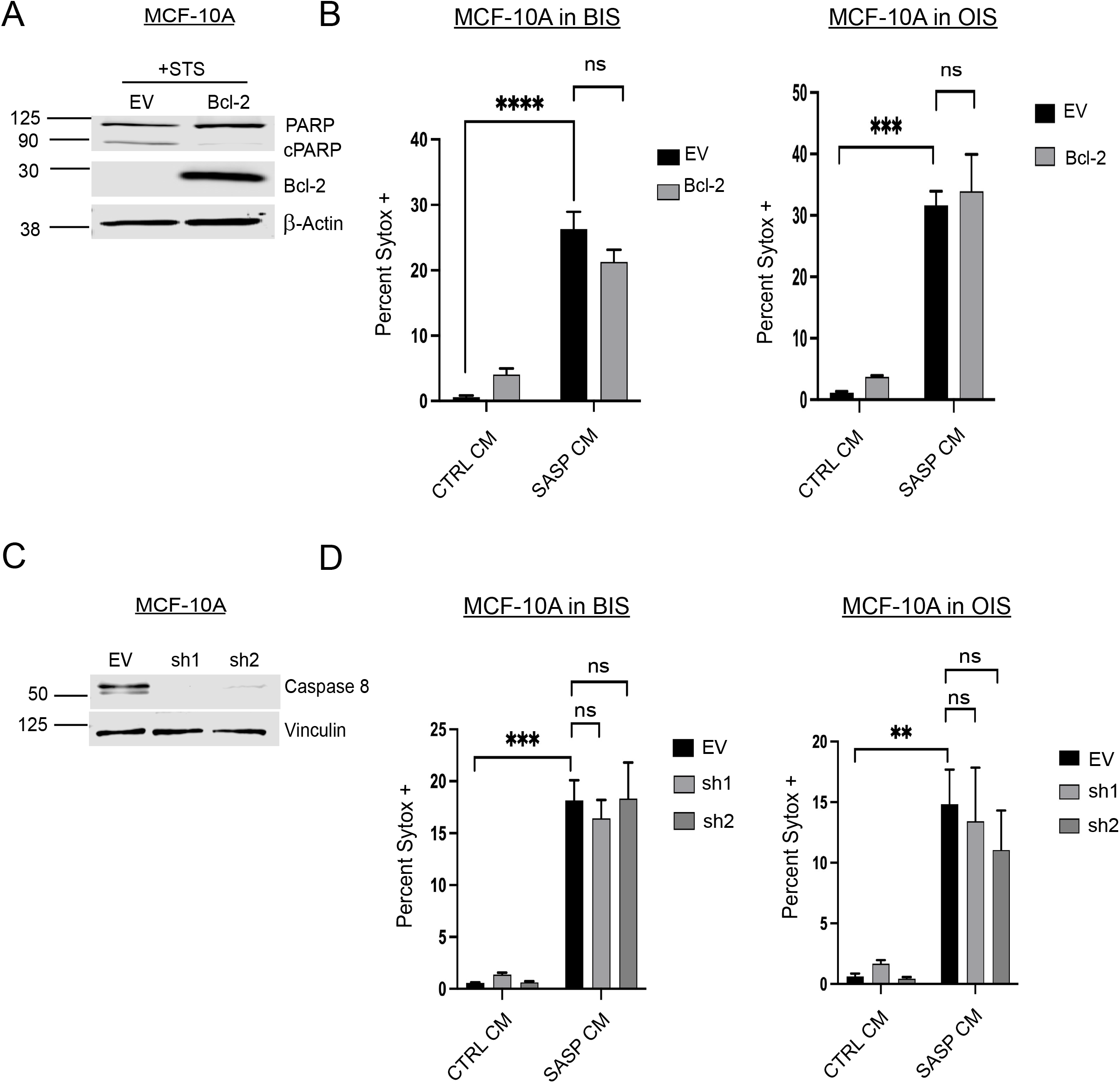
SASP CM-induced cell death is independent of the intrinsic and extrinsic apoptotic pathways. **(A)** MCF-10A cells were engineered to express EV or Bcl-2. 10A-EV and 10A-Bcl-2 cell lysates were immunoblotted to validate Bcl-2 overexpression. To confirm that the Bcl-2 overexpression is functional, both 10A-EV and 10A-Bcl-2 cells were treated with the staurosporine (STS) and probed for the cleavage of PARP, a marker of apoptosis. **(B)** 10A-EV and 10A-Bcl-2 cells were treated with CTRL or SASP CM collected from BIS or OIS fibroblasts were imaged using immunofluorescence microscopy. Cells were stained with Hoechst and SYTOX Green. Quantification of the percentage of SYTOX Green positive cells out of the total number of cells was depicted. **(C)** MCF-10A cells were engineered with shRNA against caspase-8 and immunoblotted to validate the reduction in caspase-8 protein. **(D)** Control or caspase-8 shRNA transduced cells were treated with CTRL or SASP CM from BIS or OIS and imaged using immunofluorescence microscopy. Cells were stained with Hoechst and SYTOX Green. Quantification of the percentage of SYTOX Green positive cells out of the total number of cells was depicted. SYTOX data were analyzed by two-way ANOVA followed by Tukey comparison test where ns is no statistical significance, * p<.05, ** p<0.01, *** p<0.001 and **** p<0.0001. Data are presented as mean +/− SEM. Graphs are representative data collected from a minimum of three biological replicates.

### NLRP3, Caspase-1, and Gasdermin D mediated pyroptosis underlies the capacity of SASP CM to induce cell death

Thus far, our data suggest that BIS and OIS in fibroblasts cause these cells to secrete factors that cause a caspase-dependent, but non-apoptotic cell death. When considering these surprising results, we hypothesized that the ability of z-VAD-fmk to block SASP CM-mediated cell death may be linked to its capacity to inhibit inflammatory, rather than apoptotic, caspases (47–49). Indeed, inflammatory caspases such as caspase-1 can cause cell death by pyroptosis, which is ultimately characterized by downstream activation of gasdermin proteins that form pores in the plasma membrane (50,51). In addition, the composition of the SASP includes several factors that are linked to inflammatory caspase activation through regulation of (or by) inflammasome complexes (52,53). In particular, the Nod-like receptor protein 3 (NLRP3) inflammasome can be primed or indirectly activated by inflammatory cytokines (e.g. IL-1β, IL-6) that are often found as components of the SASP (38,54–56).

In order to determine if SASP CM was causing cell death by pyroptosis in non-cancerous mammary epithelial cells, we first treated MCF-10A cells with MCC950, a small molecule inhibitor of NLRP3, and assessed the capacity of SASP CM to induce cell death. Indeed, MCC950 treatment abrogated SASP CM (from BIS or OIS) induced cell death (Figure 5A). Given these data, we next sought to ascertain if caspase-1 was required for SASP CM to induce cell death. In order to do so, we engineered MCF-10A cells to be deficient in caspase-1 levels (Figure 5B) and assessed the capacity of SASP CM to induce cell death. Our data revealed that shRNA-mediated reduction in caspase-1 significantly attenuated the induction of cell death from exposure to SASP CM (from BIS or OIS) (Figure 5C). Lastly, we assessed the importance of gasdermin D (GSDMD) in mediating SASP CM-induced cell death. During pyroptosis, the N-terminal domain of GSDMD (GSDMD-N) is separated from its C-terminal domain by caspase-1 mediated cleavage, which subsequently removes the intramolecular inhibition of the pore forming capabilities of the N-terminal domain (51). As a result, N-terminal GSDMD can permeabilize the plasma membrane to cause pyroptosis. As such, we engineered MCF-10A cells with shRNA-mediated reduction of GSDMD levels (Figure 5D) and assessed the capacity of SASP CM to promote cell death. Indeed, SASP CM was unable to promote death in cells deficient in GSDMD (Figure 5E). Taken together, our data demonstrate that the ability of SASP CM from senescent fibroblasts to promote cell death is dependent on NLRP3-mediated activation of caspase-1 and GSDMD-mediated pyroptosis.

**Figure 5:**
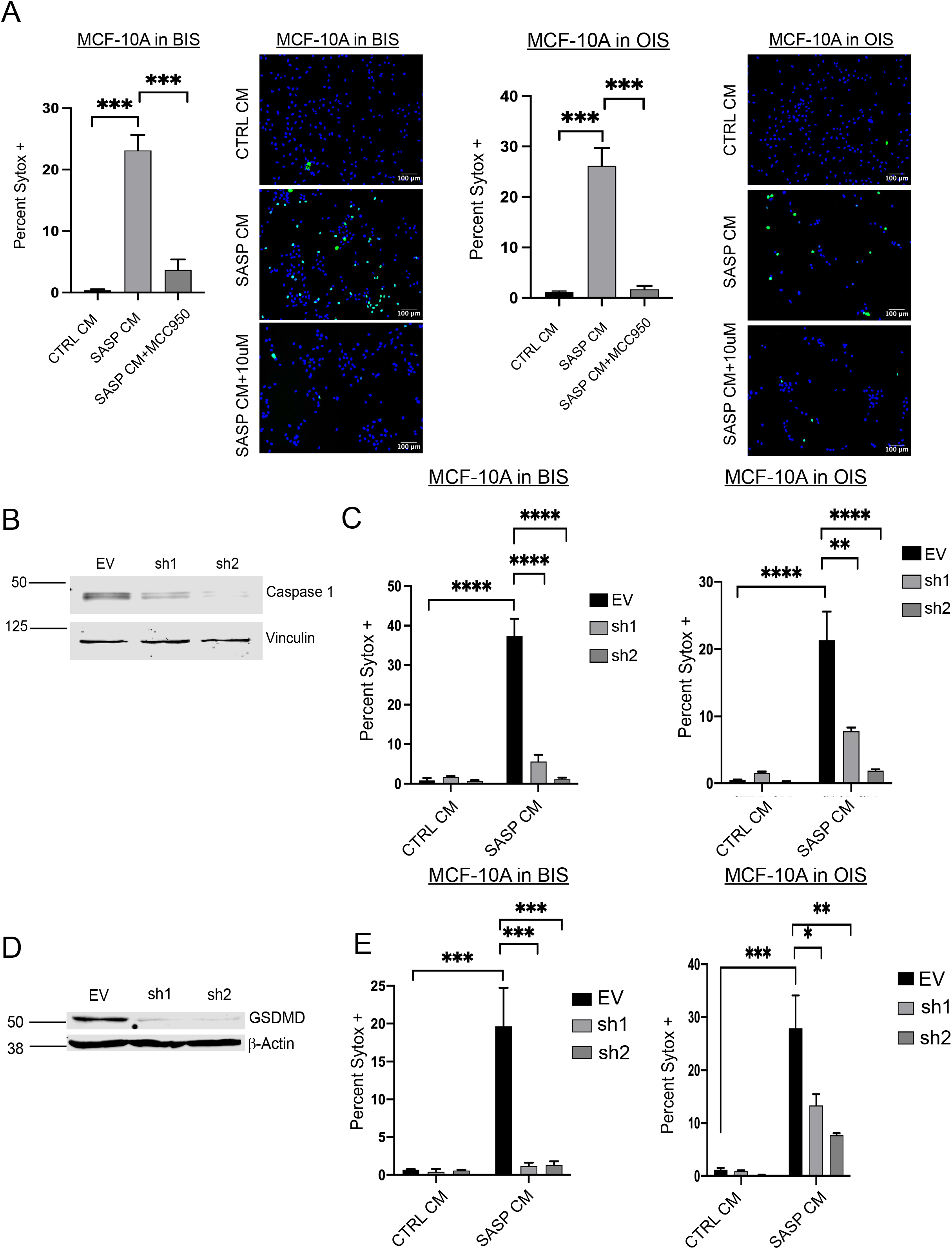
SASP CM causes cell death by pyroptosis. **(A)** MCF-10A cells were treated with CTRL or SASP CM collected from BIS or OIS fibroblasts with or without MCC950 (10uM), an NLRP3 inhibitor. Cells were imaged using immunofluorescence microscopy. Images were taken at 10X and the scale bar is 100μm. Cells were stained with Hoechst and SYTOX Green. Quantification of the percentage of SYTOX Green positive cells out of the total number of cells was depicted. **(B)** MCF-10A cells were engineered using shRNA against caspase-1 and immunoblotted to validate the reduction in caspase-1 protein. **(C)** EV Control or caspase-1 shRNA transduced cells were treated with CTRL or SASP CM from BIS or OIS and imaged using immunofluorescence microscopy. Cells were stained with Hoechst and SYTOX Green. Quantification of the percentage of SYTOX Green positive cells out of the total number of cells was depicted. **(D)** MCF-10A cells were engineered using shRNA against GSDMD and immunoblotted to validate the reduction in GSDMD protein. **(E)** EV Control or GSDMD shRNA transduced cells were treated with CTRL or SASP CM from BIS or OIS and imaged using immunofluorescence microscopy. Cells were stained with Hoechst and SYTOX Green. Quantification of the percentage of SYTOX Green positive cells out of the total number of cells was depicted. Sytox data was analyzed by two-way ANOVA followed by Tukey comparison test where ns is no statistical significance, * p<.05, ** p<0.01, *** p<0.001 and **** p<0.0001. Data are presented as mean +/− SEM. Graphs are representative data collected from a minimum of three biological replicates.

## Discussion

Our studies reveal that the induction of senescence in fibroblasts (by BIS and OIS) alter the fibroblast secretome in a fashion that promotes the induction of cell death in normal, non-transformed mammary epithelial cells. In contrast, SASP CM is unable to cause cell death in breast cancer cells or in mammary epithelial cells that have been engineered to express activated oncogenes. Furthermore, the observed cell death in normal mammary epithelial cells is blocked by z-VAD-fmk treatment, suggesting that it is dependent on the activation of caspases. Surprisingly, neither the intrinsic nor extrinsic apoptotic pathways are required for SASP CM-induced cell death. Instead, the induction of cell death is dependent on NLRP3-mediated activation of caspase-1 and the downstream actions of GSDMD. Taken together, these findings suggest that SASP CM is capable of inducing cell death by pyroptosis in normal mammary epithelial cells.

The fact that senescent fibroblasts can secrete factors that cause the induction of pyroptosis in nearby mammary epithelial cells is surprising and raises several important and interesting questions. Perhaps most salient is the ramifications in the tissue environment after the induction of a pyroptotic death, which is widely characterized to result in an inflammatory response (50). As such, the consequences of a potential influx of inflammatory cells on the remaining mammary epithelial cells and on the tissue microenvironment are potentially profound. Indeed, even if pyroptosis functions as potential mechanism to eliminate incipient neoplastic cells, the resulting inflammatory response may counterintuitively create a pro-tumorigenic niche that favors oncogenic transformation (23). Relatedly, pyroptosis has perhaps been most often studied in the context of host immune response against infections (57–60). The role of pyroptosis in cancer is less well understood, although evidence for pyroptosis playing a tumor suppressive role is accumulating in melanomas and breast cancers (61–65). Thus, it will be important to better understand the role of SASP CM-mediated pyroptosis in additional cell types and disease contexts to better appreciate resulting physiological and pathological conditions.

In addition, these surprising findings raise questions about the ramifications of therapeutic strategies that focus on the elimination of senescent cells. Given that the expression of oncogenes in normal mammary epithelial cells can prevent SASP CM-mediated cell death, it seems plausible that senescent fibroblasts are functioning in a fashion to eliminate cells that could become cancerous. Thus, senolytic agents may, in fact, function to eliminate a possible mechanism of tumor suppression. More broadly, it is unclear if senescent fibroblasts may affect adjacent epithelial cells in other tissue settings. While mammary epithelial cells respond to SASP CM by undergoing pyroptosis, it remains possible that epithelial cells from other tissues may respond in a different fashion. Moreover, additional studies investigating the anatomical location and appearance of senescent fibroblasts will help contextualize these findings to better consider the potential impacts of systemic administration of senolytic agents.

## Experimental procedures

### Cell Culture

MCF-10A cells (ATCC) and derivatives were cultured in Dulbecco’s Modified Eagle Medium Nutrient Mixture F-12 Ham 1:1 powder (Gibco) supplemented with 5% horse serum (Invitrogen), 20 ng/ml epidermal growth factor (EGF), 10 μg/ml insulin, 500 μg/ml hydrocortisone, 100 ng/ml cholera toxin, 1% penicillin/streptomycin and Plasmocin^®^ prophylactic. BJ human foreskin fibroblasts, MDA-MB-231 and MDA-MB-436 were cultured in Dulbecco’s Modified Eagle Media high glucose (DMEM) supplemented with 10% fetal bovine serum (FBS) and 1% penicillin/streptomycin. HMEC human mammary epithelial cells and derivatives (Lonza) were cultured in mammary epithelium basal medium MEBM ^TM^ (Lonza) plus MEBM SingleQuot™ Supplements. Komen Tissue bank cells (KTB-34) were cultured as previously described (66). Mycoplasma testing was routinely performed using mycoplasma PCR detection kit (LiliF, 102407-870) in all cell lines.

### Reagents

z-VAD-fmk (ApexBio) was dissolved in DMSO and used at a final concentration of 20 μM. Staurosporine (Sigma Aldrich) was dissolved in DMSO and used at a final concentration of 1μM. MCC950 (Sigma Aldrich) was dissolved in DMSO and used at a final concentration of 10μM. Sytox Green (Life Technologies) was used at a final concentration of 500nM. Hoechst 33342 was dissolved in DMSO and used at a final concentration of 0.33μg/ml. Bleomycin sulfate (ApexBio) was dissolved in DMSO and used at a final concentration of 75μg/ml. 5-Bromo-4-chloro-3-indolyl β-D-galactopyranoside (Sigma) was used at a final concentration of 1mg/ml. Senescence associated β-Galactosidase staining solution (SA-Gal) was made as previously described (67). Puromycin (Invivogen) was used at a final concentration of 1μg/ml.

### Cell Viability Assay

Cells were plated at a density of 30,000 cells per well in triplicate in 12 well plates and allowed to acclimate for 24hrs. Cells were treated with either CTRL or SASP CM for an additional 24hrs prior to imaging. Cells were stained first in Hoechst 33342 in cell culture media for 15 minutes protected from light at a final concentration of 0.33μg/ml. Hoechst stain was removed and cells were washed in 1X PBS. Cells were stained in culture media plus SYTOX Green at a final concentration of 500nM for 15 minutes and protected from light. The SYTOX Green stain was removed, cells were washed in 1X PBS and then placed back in normal culture medium while images were taken using Zeiss Axio Observer.A1 inverted fluorescent microscope using ZEN 2012 software. Images were taken in brightfield using the 10X objective. Images were merged and quantified using the cell counter plug in on FIJI (National Institutes of Health, Bethesda, MD, USA). Five images were taken per well across three wells per condition. Averages, standard deviation and SEM were calculated. Data are represented as average percentage SYTOX out of the total number of cells. Representative data from at least three biological replicates are shown.

### Western Blot Analysis

After treatment, cells were plated in a 6 well plate at a density of 200,000 cells and whole cell lysates were collected at 24hrs (68–70). Cells were washed one time with 2mL PBS and lysed in RIPA buffer (20mM Tris-HCl (pH 7.5), 150 mM NaCl, 1 mM Na2EDTA, 1 mM EGTA, 1% NP-40, 1% sodium deoxycholate) supplemented with 1 μg/ml aprotinin, 5 μg/ml leupeptin, 20 μg/ml phenylmethylsulfonyl fluoride (PMSF) and HALT phosphatase inhibitor mixture (Thermo Scientific). Lysis buffer is added directly to the wells and the cells are lysed on the plate. Cell lysates are sonicated with 1-s pulse on and 1-s pulse off for 8 s using Branson sonifier at 10% amplitude. After sonication, cell lysates were cleared by centrifuging at 13,000 rpm for 15 min. Supernatants were quantified by BCA assay (Pierce Biotechnology, Waltham, MA, USA) and normalized to the same protein concentrations. 6X Lamelli sample buffer was added to the cell lysates, and they were boiled on a heat block at 95°C for 10 min. After running samples on SDS-PAGE alongside Chameleon Duo Pre-stained protein ladder (LI-COR), gels were then transferred onto Immobilon-FL PVDF membrane (Millipore) and blocked for 45 minutes in 5% milk. Membranes were probed in 2% milk along with primary antibodies at 4°C overnight. The following antibodies were used for western blotting: ß-Actin (Sigma-Aldrich; no. A1978), Caspase 8 (Proteintech; 13423-1-AP), Caspase 1 (Proteintech; 22915-1-AP), GSDMD (CST;69469) LaminB1(Proteintech; 66095-1), RAS (CST; 3965S), PARP/cPARP (CST; 9542S), Bcl-2; Proteintech (12789-1-AP), Vinculin (Proteintech; 66305-1-Ig), HER2 (Cell Signaling Technology; 2242S). Donkey anti-Mouse IgG (H+L) Alexa Fluor™ Plus 680 Highly Cross-Adsorbed Secondary Antibody and Donkey anti-Rabbit IgG (H+L) Alexa Fluor™ Plus 800 Highly Cross-Adsorbed Secondary Antibody were incubated for 1hr in 2% milk protected from light. Membranes were imaged using Odyssey CL_X_ Infared Imaging system by LI-COR Biosciences and analyzed using Image Studio Version 5.2.

### Generation of stable cell lines using retrovirus

Generation of stable cell lines were made using the pBABE-Puro-based retroviral vectors encoding the overexpression of HRAS, Bcl-2 and ErbB2 were used to generate stable cell lines. HEK293T cells were transfected with target DNA (10ug) along with the packaging vector pCLAmpho (10ug) using the Calcium phosphate transduction method. This method uses 2X HBSS (280 mM NaCl, 10 mM KCl, 1.5 mM Na_2_HPO_4_, 12 mM Glucose, 50 mM HEPES, pH 7.05) and 2.5M CaCl_2_ and sterile water. Supernatants were collected 48-hours posttransfection, filtered through a 0.45μm filter (EMD Millipore), and used for viral transduction. Stable populations of puromycin-resistant cells were obtained by selecting in 1ug/mL puromycin (Invivogen, San Diego, CA, USA) for at least 48hrs.

### Generation of stable knockdown cell lines using lentiviral delivery of shRNA

MISSION short hairpin RNA (shRNA) constructs against CASP8 (NM 033356 clone TRCN0000376482 and TRCN0000377309) in the puromycin-resistant pLKO.4 vector along with an empty vector control were purchased from Sigma-Aldrich. MISSION short hairpin RNA (shRNA) constructs against CASP1 (NM_001223 clones TRCN0000003502 and TRCN00000010796) in the puromycin-resistant pLKO.4 vector along with an empty vector control were purchased from Sigma-Aldrich. MISSION short hairpin RNA (shRNA) constructs against GSDMD (NM_024736.4 clones TRCN000000179394 and TRCN000000180013) in the puromycin-resistant pLKO.4 vector along with an empty vector control were purchased from Sigma-Aldrich. For packaging of lentivirus, HEK293T cells were transfected with 2-3ug target DNA along with the packaging vectors psPAX2 (200ng) and pCMV-VSV-G (300ng) using Lipofectamine 2000. Virus was collected 48-hours post-transfection, filtered through a 0.45μm filter (EMD Millipore), and used for infecting MCF-10A, HMEC and KTB-34 cells in the presence of 10μg/ml polybrene. Stable populations of cells were selected using 1μg/mL puromycin for at least 48h. Immunoblots were performed to confirm success of knockdowns.

### RNA isolation and quantitative real-time PCR

Total RNA was isolated with Zymo Research Quick-RNA^TM^ Mini Prep RNA kit. Cells were lysed directly on the plate. Total mRNA was extracted according to manufacturer’s protocol. Reverse transcription was performed using 1μg of RNA using an iScript Reverse Transcription Supermix (Bio-Rad, 1708840) to generate cDNA. This cDNA was utilized for qRT-PCR using SYBR green supermix (Bio-RAD, 1725271) and specific primers on a 7,500 fast real-time PCR system (Applied Biosystems, Life Technologies, Waltham, MA, USA). The relative levels of gene transcripts were normalized to GAPDH which were determined by quantitative real-time PCR. The primer sequence for specific transcripts were as followed: Human GAPDH Forward: 5’-GCATGGCCTTCGGTGTCC, Human GAPDH Reverse: 5’-AATGCCAGCCCCAGCGTC-AAA, Human IL-6 Forward: 5’ACATCCTCGACGGCATCT-CA, Human IL-6 Reverse: 5’-TCACCAGGCAAGTCTCCT-CA, Human IL-8 Forward: 5’GCTCTGTGTGAAGGTGCAGT, Human IL-8 Reverse: 5’-TGCACCCAGTTTTCCTTG-GG, Human CCL20 Forward: 5’-CTGCGGCGAATCAGAAGC-AGC. Human CCL20 Reverse: 5’-CCTTCATTGGCCAGCTGC-CGT. Amplification was carried out at 95C for 12 min, followed by 40 cycles of 15 s at 95C and 1 minute at 60C. Standard error of the mean is represented by the error bars and P-values were calculated using a two-tailed t test.

### Conditioned media generation

Cells were seeded at 90,000 cells for DMSO treatment and 200,000 cells for Bleomycin treatment respectively onto a 10cm^2^ plate and allowed to grow 48hrs prior to senescence induction using Bleomycin or control treatment with DMSO. Cells were treated for 48h, media was removed, and cells were washed with 1X PBS two times and placed in DMEM-F12 media to condition for a period of 7 days for senescence and 1 day for control plates. For the onocogene-induced senescence model, retrovirus was used to create pBABE-EV and pBABE-HRas(G12V) virus. Fibroblasts were infected for 24hrs with supernatant, selected in 1μg/mL Puromycin for 48hrs. EV CTRL cells were conditioned in DMEM-F12 media for 24hrs and HRAS cells were conditioned for 7 days as described above. Media was collected and filtered through 0.2μm PVDF filter.

### SA β-Gal Staining

Cells were fixed using 4% PFA diluted 1:1 with 1X PBS for 5 minutes. Cells were then washed three times using 1X PBS and stained using the chromogenic assay as previously described (67). Images were captured on an Axio Observer Inverted microscope using Zen 2.5 (blue edition) software for image processing. All images were taken at 10X in brightfield.

### Quantification and Statistical Analysis

As indicated in the legends, quantitative data are represented as the mean+/− standard error of the mean (SEM). All graphs and statistical analysis were performed using Graphpad Prism (9.0.0). Statistical significance with two groups was calculated using a student two-tailed t test. Each graph represents data collected from a minimum of three biological replicates. Group comparisons were made using one or two-way analysis of variance (ANOVA) followed by Tukey test.

## Data availability

Further information and requests for resources, reagents, or data will be fulfilled by the corresponding author Zachary T. Schafer.

## Acknowledgements

We thank Veronica Schafer and all current/past Schafer lab members for helpful comments and/or valuable discussion. We thank Harikrishna Nakshatri (Indiana University School of Medicine) for the KTB-37 cell line, Ana Flores-Mireles (University of Notre Dame) for experimental assistance, Athanasia Panopoulos (Cedars-Sinai) for the BJ cell line, and the Notre Dame Integrated Imaging Facility for assistance with microscopy.

## Author contributions

LMH, SS, JC, PL, SC, and IMM conducted experiments, analyzed data, and interpreted results. LMH and ZTS wrote the manuscript with feedback from all other authors. LMH and ZTS were responsible for conception/design of the project. ZTS was responsible for overall study supervision.

## Funding and additional information

ZTS is supported by the National Institutes of Health/National Cancer Institute (R01CA262439), the Coleman Foundation, the Malanga Family Excellence Fund for Cancer Research at Notre Dame, the College of Science at Notre Dame, the Department of Biological Sciences at Notre Dame, and funds from Mr. Nick L. Petroni. This content is solely the responsibility of the authors and does not necessarily represent the official views of the National Institutes of Health.

## Conflict of interest

The authors declare that they have no conflicts of interest with the contents of this article.

## Notes

### Competing Interest Statement

The authors have declared no competing interest.

